# Characteristics of reproductive tract microbiota in health and disease

**DOI:** 10.64898/2025.12.20.695662

**Authors:** Qiang Han, Nan Wang, Teng Feng, Yue-Yue Li, Zhuang Zhu, Teng-Lin Xu, Ke-Ji Quan, George Fei Zhang

## Abstract

The symbiotic relationship between the microbiota and the host is crucial to host health. The reproductive tract microbiota in health and disease was examined by next-generation sequencing for the first time in this study. A total of 10 healthy and 30 diseased samples were collected for microbiota analysis. Species distributions, alpha diversity, beta diversity, and functional gene prediction were analyzed. *Riemerella anatipestifer*, *Escherichia coli*, *Avibacterium paragallinarum*, *Ornithobacterium rhinotracheale*, *Enterococcus faecalis*, and *Acinetobacter baumannii* were first identified as pathogenic bacteria at higher abundance in the reproductive tract of the laying hens. The diseased group exhibited dysbacteriosis and a different species-abundance cluster compared with the healthy control. Alpha-diversity analysis showed that the ACE index in the control group was significantly higher than in the diseased group. Beta diversity analysis indicates that samples from the healthy group had a more similar composition than those from the diseased group. Moreover, functional gene prediction analysis indicated that diseased groups showed a high level of gram-negative, facultative anaerobic, and potentially pathogenic bacteria. The pathogenic bacteria identified in this study are helpful for vaccine development and disease treatment. In conclusion, next-generation sequencing is an effective method for analyzing bacterial communities to support health assessment and disease diagnosis.

## 1 Introduction

Salpingitis is a significant disease of laying hens that seriously affects the profitability of chicken farms. Polymerase chain reaction (PCR) is a widely used method for detecting pathogenic agents in disease diagnosis. To protect animals from infectious disease, vaccines are a valuable tool for preventing infections caused by pathogens ^1,2^. Pathogenic infections in both the intestine and the oviduct lead to dysfunction of egg production in laying hens through inflammation induced by mucosal immune responses ^3^.

*Salmonella enteritidis* could be detected in oviduct tissues associated with egg formation ^4,5^. The ratio of isolated bacteria per isolated tubular gland cell was again significantly higher in the isthmus as compared with in the magnum. Tubular gland cells of the isthmus were more heavily invaded than those of the magnum ^6^. The role of type 1 fimbriae in the interaction of *Salmonella enteritidis* with the hen’s oviduct was well studied. *Salmonella enteritidis* adheres to the surface of the epithelium of the chicken oviduct by type 1 fimbriae ^7^. *Salmonella enteritidis* genes were induced during oviduct colonization and egg contamination in laying hens ^8^.

Chicken oviduct epithelial cells express most of the known AvBD genes in response to SE infection. PipB, a T3SS-2 effector protein, plays a role in dampening the β-defensin arm of innate immunity during SE invasion of chicken oviduct epithelium ^9^. *E. coli*-associated egg peritonitis was responsible for 15.39% of the reproductive tract abnormalities in commercial layers between 21 and 80 week of age ^10^. *Gallibacterium anatis* (*G. anatis)* has been implicated in salpingitis and peritonitis in egg-laying chickens, leading to decreased egg production and increased mortality worldwide ^11^. This view was substantiated by Mirle et al. (1991), who, in a postmortem investigation including 496 hens, isolated *Gallibacterium* as one of the most frequent bacterial agents from lesions in the reproductive tract ^12^. *Salmonella Enteritidis* downregulates flagellar gene expression in the oviduct and consequently prevents a flagellin-induced inflammatory response, thereby increasing its oviduct colonization efficiency ^13^.

16S rRNA sequencing is a valuable tool to analyze microbial communities ^14,15^. In this study, the chicken reproductive tract was analyzed using 16S rRNA sequencing. Healthy and diseased reproductive tracts were analyzed for microbial community, alpha diversity, beta diversity, and biomarkers. This study aims to identify a novel pathogenic agent of chicken salpingitis.

## 2 Materials and Methods

### 2.1 Sample collection and study design

Samples were collected from chickens with salpingitis for microbial community analysis. Samples were collected from SPF chicken oviducts as a negative control. Ten negative controls and thirty diseased samples were included in this study (Table 1). Wohua Bioeng Experimental Animal Center approved animal experiments. All the samples were collected for 16S rRNA sequencing.

**Table 1.** Samples collected in this study.

| Number | Breed | Day | Source | Symptom |
| --- | --- | --- | --- | --- |
| CK-1 | Leghorn | 120 | Binzhou/Shandong | - |
| CK-2 | Leghorn | 120 | Binzhou/Shandong | - |
| CK-3 | Leghorn | 120 | Binzhou/Shandong | - |
| CK-4 | Leghorn | 120 | Binzhou/Shandong | - |
| CK-5 | Leghorn | 120 | Binzhou/Shandong | - |
| CK-6 | Leghorn | 120 | Binzhou/Shandong | - |
| CK-7 | Leghorn | 120 | Binzhou/Shandong | - |
| CK-8 | Leghorn | 120 | Binzhou/Shandong | - |
| CK-9 | Leghorn | 120 | Binzhou/Shandong | - |
| CK-10 | Leghorn | 120 | Binzhou/Shandong | - |
| Oviduct-1 | Hy-Line | 167 | Rizhao/Shandong | Swollen head syndrome |
| Oviduct-2 | Hy-Line | 160 | Qingdao/Shandong | Swollen head syndrome |
| Oviduct-3 | Hy-Line | 63 | Dezhou/Shandong | Lethargy and decreased appetite |
| Oviduct-4 | Hy-Line | 312 | Heze/Shandong | Oviductitis |
| Oviduct-5 | Hy-Line | 462 | Sichuan | Oviductitis |
| Oviduct-6 | Hy-Line | 347 | Haerbin/Heilongjiang | Diarrhoea |
| Oviduct-7 | Hy-Line | 221 | Rizhao/Shandong | Lethargy and decreased appetite |
| Oviduct-8 | Hy-Line | 160 | Binzhou/Shandong | Swollen head syndrome |
| Oviduct-9 | Hy-Line | 417 | Qingtongxia/Ningxia | Abnormal eggs, diarrhoea |
| Oviduct-10 | Hy-Line | 150 | Hengshui/Hebei | Lethargy and decreased appetite |
| Oviduct-11 | Hy-Line | 200 | Dezhou/Shandong | Lethargy and decreased appetite |
| Oviduct-12 | Hy-Line | 190 | Qichun/Hubei | Lethargy and decreased appetite |
| Oviduct-13 | Hy-Line | 115 | Liaocheng/Shandong | Diarrhoea |
| Oviduct-14 | Hy-Line | 172 | Weifang/Shandong | glandular gastritis, necrotic enteritis, peritonitis |
| Oviduct-15 | Hy-Line | 180 | Baotou/Neimeng | Oviductitis, |
| Oviduct-16 | Hy-Line | 40 | Binzhou/Shandong | Lethargy and decreased appetite |
| Oviduct-17 | Hy-Line | 160 | Wuhan/Hubei | Oviductitis |
| Oviduct-18 | Hy-Line | 331 | Binzhou/Shandong | Sequencing failed |
| Oviduct-19 | Hy-Line | 95 | Suzhou/Anhui | Oviductitis |
| Oviduct-20 | Hy-Line | 170 | Linyi/Shandong | diarrhoea, breathlessness |
| Oviduct-21 | Hy-Line | 309 | Dezhou/Shandong | Oviductitis |
| Oviduct-22 | Hy-Line | 314 | Dezhou/Shandong | Oviductitis |
| Oviduct-23 | Hy-Line | 378 | Rizhao/Shandong | Oviductitis |
| Oviduct-24 | Hy-Line | 372 | Dezhou/Shandong | Reduced egg production |
| Oviduct-25 | Hy-Line | 391 | Dezhou/Shandong | Oviductitis |
| Oviduct-26 | Hy-Line | 328 | Rizhao/Shandong | Oviductitis |
| Oviduct-27 | Hy-Line | 322 | Rizhao/Shandong | Oviductitis |
| Oviduct-28 | Hy-Line | 381 | Qingdao/Shandong | Oviductitis |
| Oviduct-29 | Hy-Line | 140 | Liaocheng/Shandong | Swollen head syndrome, Oviductitis |
| Oviduct-30 | Hy-Line | 265 | Yuncheng/Shanxi | Swollen head syndrome, Oviductitis |

### 2.2 Library construction and sequencing

After extracting the total DNA of the sample, primers were designed based on conserved regions. Sequencing adapters were added to the ends of the primers, and PCR amplification was performed. The products were then purified, quantified, and homogenized to form a sequencing library. The constructed library was first subjected to library quality control. Libraries that passed quality control were sequenced on an Illumina NovaSeq 6000. The original image data files obtained by high-throughput sequencing (such as Illumina NovaSeq and other platforms) are converted into original sequencing reads (Sequenced Reads) through base-calling analysis. The results are stored in FASTQ (abbreviated as fq) file format, which contains the sequence information of the sequencing sequence (Reads) and its corresponding sequencing quality information. Amplicon 16S V3+V4 was amplified with primers (F: ACTCCTACGGGAGGCAGCA; R: GGACTACHVGGGTWTCTAAT). Species annotation was conducted using BLAST and Bayesian analysis with the Silva 138 database.

### 2.3 Data processing

Raw Reads obtained by sequencing were filtered with Trimmomatic v0.33, and cutadapt 1.9.1 was used to identify and remove primer sequences, yielding Clean Reads. Use the dada2 method in QIIME2 2020.6 to denoise, splice double-end sequences and remove chimeric sequences to obtain the final valid data (Non-chimeric Reads). Information analysis content: Feature division (OTUs, ASVs), diversity analysis, difference analysis, correlation analysis and function prediction analysis (see analysis results for details).

### 2.4 Alpha diversity analysis

Alpha diversity reflects the richness and diversity of species within a single sample, and several metrics can be used to measure it. Alpha-diversity index differences can be visualized in a box plot of the data in the table above to demonstrate differences in alpha diversity across sample groups. The significance of the difference is assessed using the Student’s t-test. The dilution curve is constructed by randomly selecting a given number of sequences from the sample, counting the number of species represented by these sequences, and plotting the sequence number against the number of species. This is used to determine whether the amount of sequencing data is sufficient to reflect the species diversity in the sample and, by extension, the species richness in the sample. The cumulative relative abundance curve of species reflects the relationship between the number of samples and the number of annotated species.

### 2.5 Beta diversity analysis

Beta diversity analysis was performed using QIIME to compare species diversity among samples. Principal Component Analysis (PCA) is a technique for analyzing and simplifying datasets by decomposing variance and representing differences between datasets on a two-dimensional coordinate plane. PCA analysis is based on a ranking analysis of the original species composition matrix.

### 2.6 Analysis of significant differences between groups

LefSe analysis, also known as the analysis of species with significant differences between groups, uses linear discriminant analysis (LDA) to estimate the magnitude of the effect of each component (species)’ abundance on the difference effect. The primary purpose of this analysis is to find species with significant differences in abundance between groups. It can identify biomarkers with statistically significant between-group differences.

### 2.7 Functional gene prediction analysis

BugBase first normalizes OTUs by the predicted 16S copy number and then predicts microbial phenotypes using a pre-computed file. BugBase selects the coverage threshold with the highest variance across all samples for each feature in the user data. BugBase generates a final biological-level trait prediction table that contains the trait’s predicted relative abundance for each sample. Predicted phenotype types include Gram-positive, Gram-negative, biofilm-forming, Pathogenic, mobile-element-containing, oxygen-utilizing, and oxidative-stress-tolerant.

## 3 Results

### 3.1 Library construction and sequencing

The sequencing of oviduct-18 failed (Table 1). A total of 2,512,282 pairs of reads were obtained from the sequencing of 39 samples. After quality control and splicing of double-end reads, a total of 2,198,741 clean reads were generated. Each sample generated at least 35,621 clean reads, with an average of 56,378.

### 3.2 Pathogenic bacteria were identified with high abundance

Using SILVA as a reference database, a Naive Bayes classifier combined with alignment was employed to annotate feature sequences, yielding species-level classification for each feature (Figure 1). *Riemerella anatipestifer*, *Escherichia coli*, *Avibacterium paragallinarum*, *Ornithobacterium rhinotracheale*, *Enterococcus faecalis*, and *Acinetobacter baumannii* were identified from animals with salpingitis (Figure 1C). Unclassified Bacteria, *Lysinibacillus fusiformis*, unclassified *Muribaculaceae*, and *Ruminococcus torques*, were identified in healthy animals (Figure 1C). Genus Riemerella, *Escherichia*_ *Shigella*, *Avibacterium*, *Ornithobacterium*, *Enterococcus*, and *Acinetobacter* were dominant in disease and *Bacteroides*, unclassified Bacteria, *Lysinibacillus*, *Exiguobacterium* are dominant in health (Figure 1B). At the family level, *Weeksellaceae*, *Enterobacteriaceae*, *Pasteurellaceae*, *Enterococcaceae,* and *Moraxellaceae* are dominant in disease and *Lachnospiraceae*, *Bacteroidaceae*, *Lactobacillaceae*, *unclassified Bacteria*, and *Planococcaceae* are dominant in health (Figure 1A).

**Figure 1.**
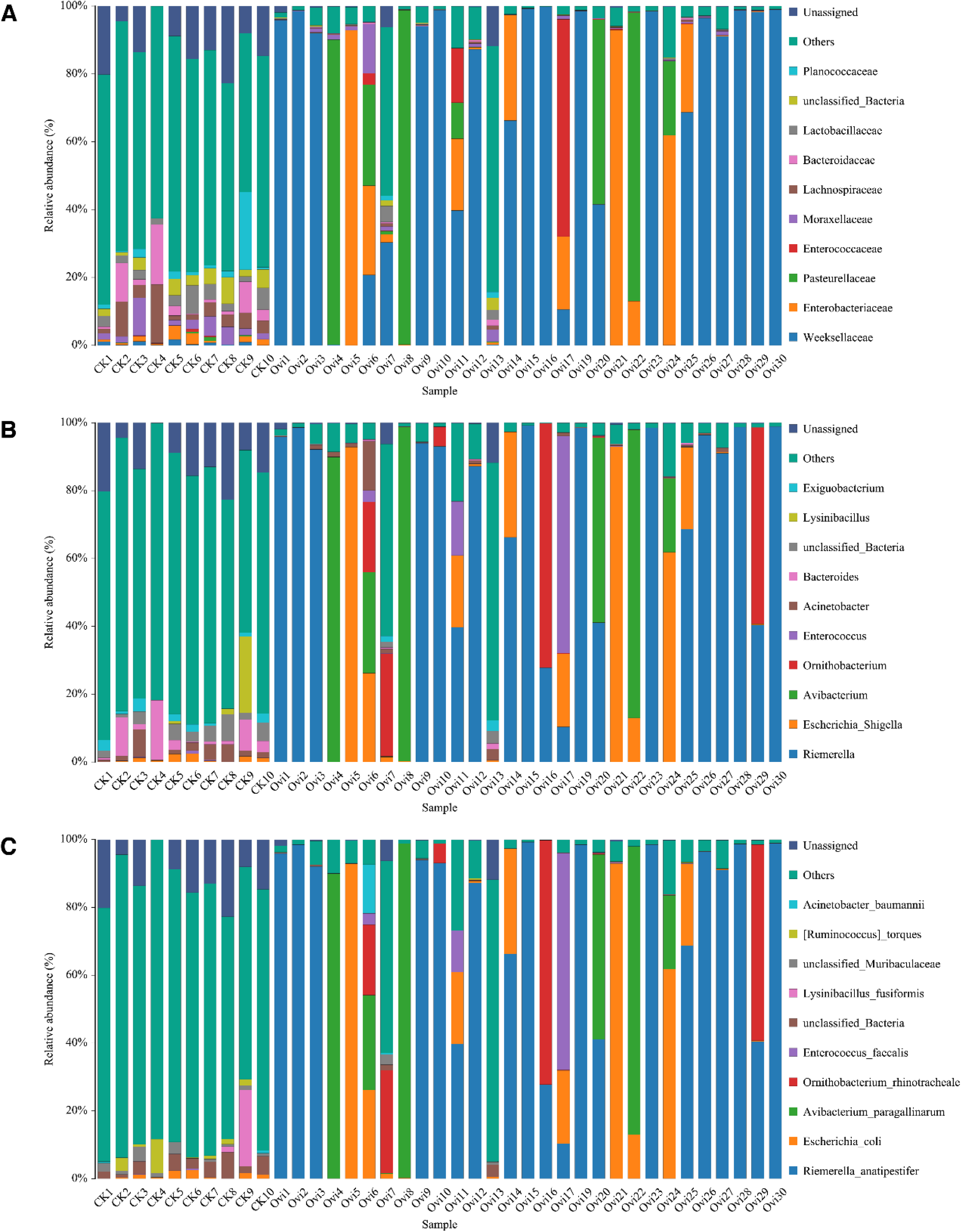
The species distribution histogram at each taxonomic level displays, from left to right, the family, genus, and species levels. Each color represents a species, and the length of the color block indicates the species’ relative abundance. The top ten species by abundance are displayed, and all other species are grouped as “Others” in the chart. Unclassified species represent unannotated taxonomies; specific species information can be found in the species abundance table at the corresponding taxonomic level. The horizontal axis indicates the sample name, and the vertical axis indicates the relative abundance percentage.

### 3.3 Species abundance cluster analysis

A heatmap is a graphical display method that uses color gradients to represent values in a data matrix and to cluster species or samples based on similarity in abundance (Figure 2). A species heatmap was constructed based on species composition and relative abundance for each sample. Heatmap cluster analysis was performed at the species level. Vertical clustering indicates similarity in species abundance across samples. The shorter the branch length, the more closely the two species are related, indicating greater similarity in species abundance across samples. Horizontal clustering indicates similarity in species abundance across samples. Similar to vertical clustering, the shorter the branch length between two samples, the greater the similarity in species abundance between them. High- and low-abundance species cluster together, and the color gradient and degree of similarity reflect differences in community composition across multiple samples. Ovi7 and Ovi13 were clustered into the control group, and most of the diseased samples from Ovi1 to Ovi30 were clustered into the same branch. *Riemerella anatipestifer*, *Acinetobacter towneri* ^16^, Rikenellaceae bacterium DTU002, unclassified Dysgonomonadaceae, Aliarcobacter cryaerophilus ^17^, Acinetobacter sp., *Flavobacterium cucumis*, *Avibacterium paragallinarum*, *Escherichia coli*, *Mycoplasmopsis synoviae*, *Ornithobacterium rhinotracheale*, *Acinetobacter baumannii*, *Enterococcus faecalis* ^18^, *Enterococcus cecorum* ^19^, *Gallibacterium anatis*, and *Actinomyces europaeus* are identified in the diseased group, which are in the same branch.

**Figure 2.**
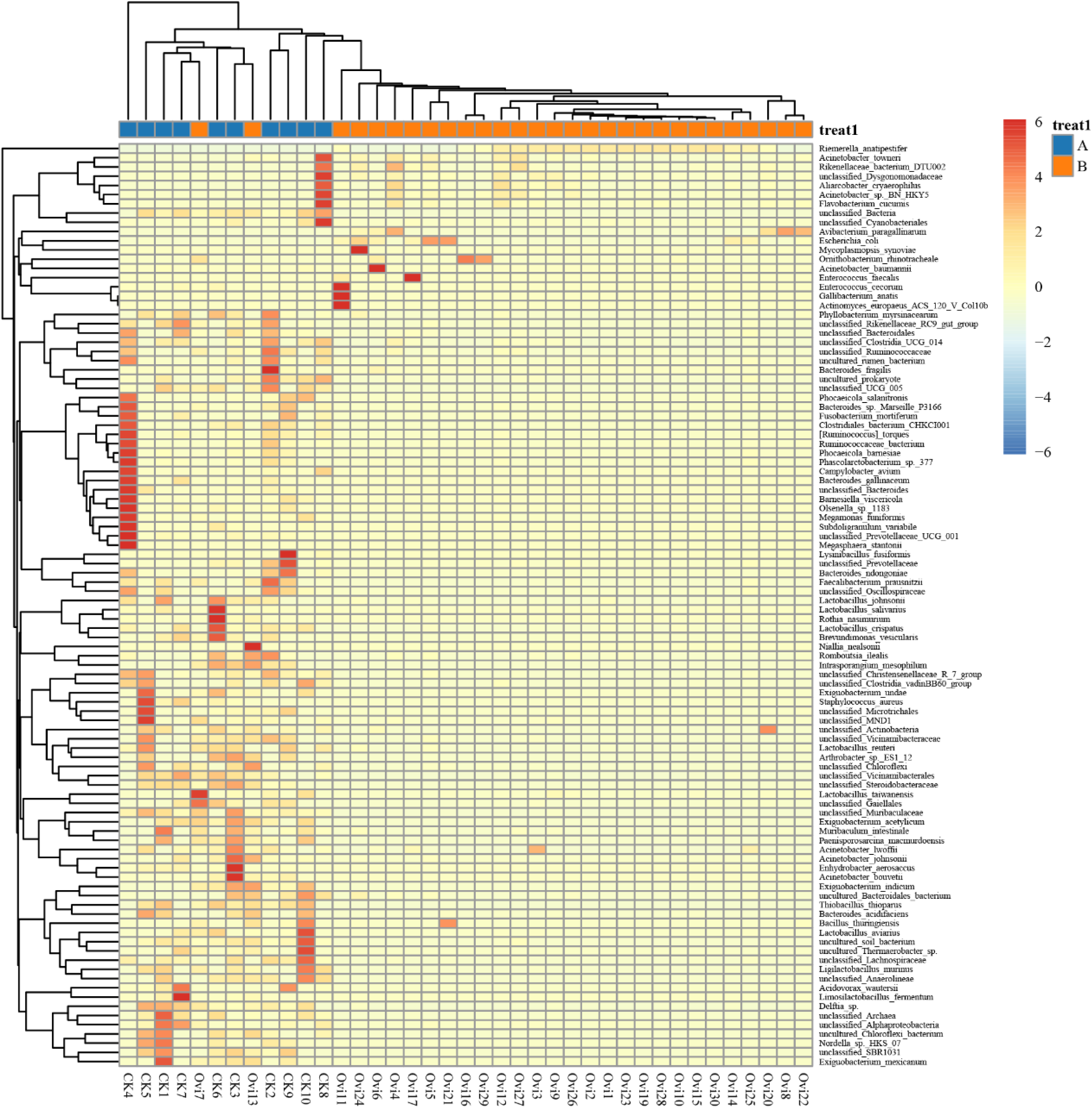
The sample species abundance cluster heat map. The result file contains cluster heat maps for each species. The values corresponding to the heat map in the figure are Z-score normalized using the R scale function for the same species across different samples. The color gradient from blue to red indicates the abundance from low to high among samples, and the colors correspond to the columns in the figure.

### 3.4 Alpha diversity analysis

Alpha diversity reflects the richness and diversity of species within a single sample. The horizontal axis represents the group name, and the vertical axis represents the corresponding alpha diversity index value (Figure 3). The numbers on the lines connecting the bars are the p-values of the t-test (Figure 3A). The ACE index in the control group is significantly higher than that in the diseased group (Figure 3A). The rarefaction curve is constructed by randomly sampling a certain number of sequences from a sample, counting the number of species represented by these sequences, and plotting the number of sequences against the number of species. It is used to assess whether the volume of sequencing data is sufficient to capture the species diversity in the sample and, indirectly, to estimate the sample’s species richness. It demonstrated that the multiple-sample/group rarefaction curves for the control were higher than those for the diseased (Figure 3B and 3C). The species relative abundance accumulation curve (Figure 3D) reflects the relationship between the number of samples and the number of annotated species.

**Figure 3.**
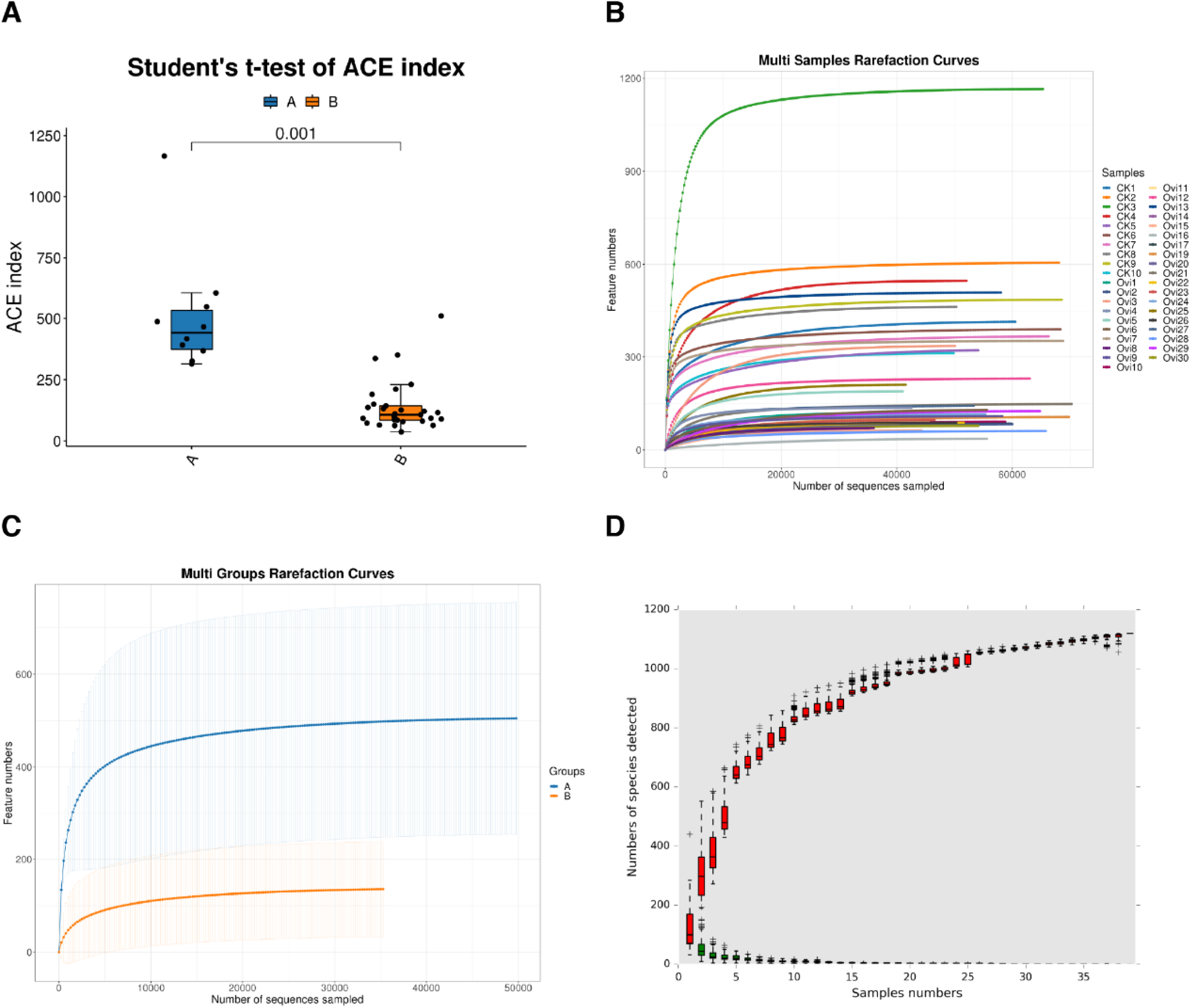
Alpha diversity analysis. (A)Alpha diversity index intergroup difference box plot. The horizontal axis shows the group name: A for the negative control and B for the clinical samples, and the vertical axis shows the corresponding alpha diversity index. (B) Genera-level species accumulation curve. (C) Multi-sample rarefaction curve. The horizontal axis represents the number of randomly selected sequencing reads, and the vertical axis represents the number of features obtained based on that number of sequencing reads. Each curve represents a sample and is colored differently. (D) Multi-group rarefaction curve. The X axis represents the number of randomly selected sequencing reads, and the Y axis represents the number of features obtained based on that number of sequencing reads.

### 3.5 Beta diversity analysis

Principal Component Analysis (PCA) was conducted to analyze beta diversity (Figure 4). The coordinate axes are defined by the two feature values that best reflect the variance. The results of the PCA analysis between groups are as follows. PCA is a dimensionality-reduction technique based on the original species composition matrix. In the figure, each point represents a sample; different colors represent different samples/groups (if applicable). PC1 (25.56%), PC2 (18.44%), and PC3 (14.17%) denote the first, second, and third principal components, and the percentages indicate each component’s contribution to the sample variance (Figure 4). The closer the two samples are, the more similar their compositions are. The elliptical circles represent 95% confidence ellipses (i.e., if there were 100 samples in that group, 95 of them would fall within the ellipse). The elliptical circles indicate that samples from the healthy group had a more similar composition than those from the diseased group (Figure 4).

**Figure 4.**
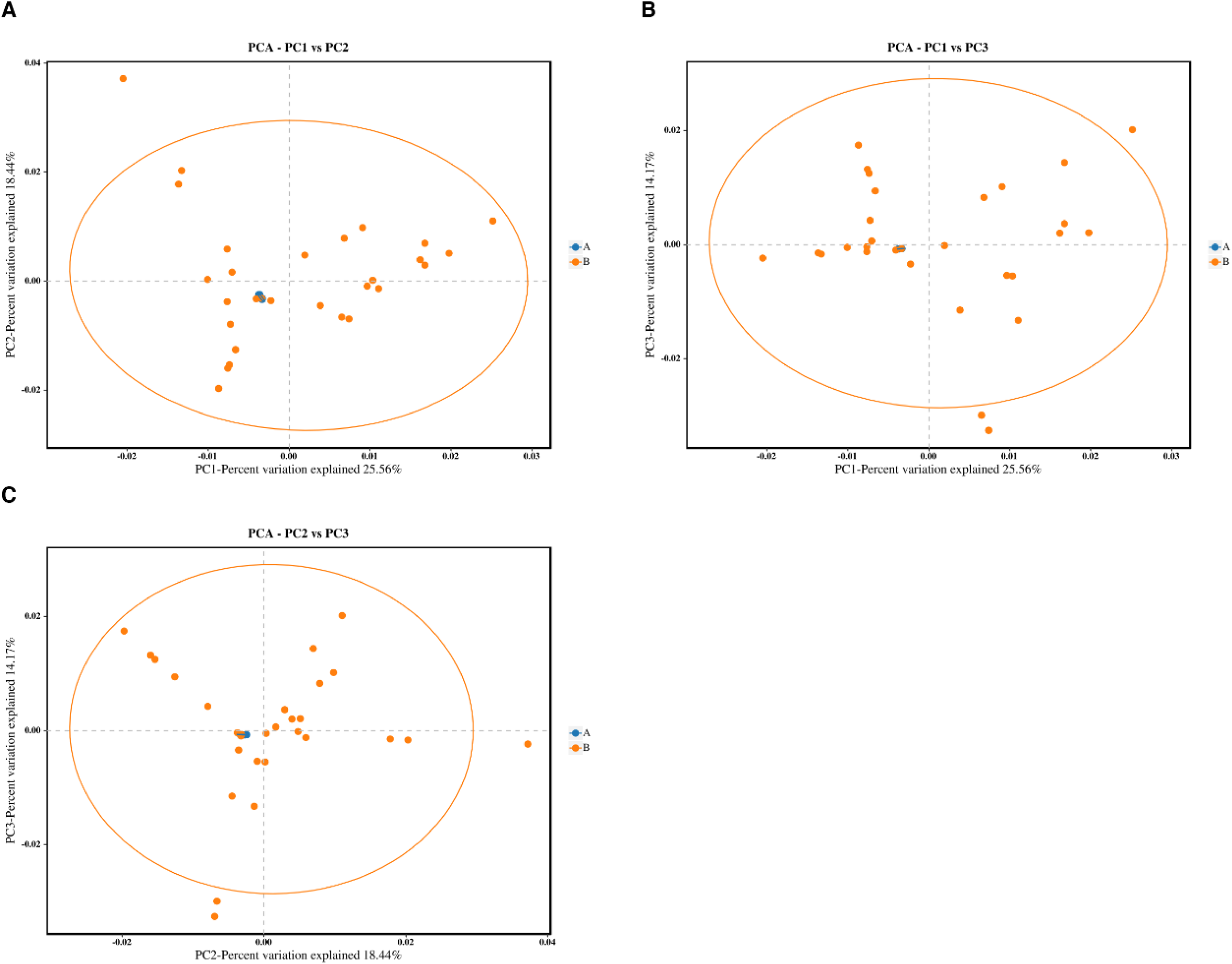
Beta diversity analysis. PCA Analysis Chart. Each point in the graph represents a sample; different colors represent different samples/groups; the ellipse indicates a 95% confidence ellipse. The X axis represents the first principal component, and the percentage represents the contribution of the first principal component to the sample differences; the Y axis represents the second principal component, and the percentage represents the contribution of the second principal component to the sample differences.

### 3.6 Analysis of significant differences between groups

LEfSe Line Discriminant Analysis (LDA) Effect Size is an analytical method that combines the non-parametric Kruskal-Wallis and Wilcoxon rank sum tests with the linear discriminant analysis (LDA) effect size (Figure 5). It can identify biomarkers with statistically significant between-group differences. Bar graph showing the abundance of biomarkers across different groups (Figure 5). *Riemerella anatipestifer* is a biomarker of the disease group, and its abundance in the diseased group is significantly higher than in the control group (Figures 5A and 5 B). Meanwhile, the abundance of *Avibacterium paragallinarum* in the disease group is significantly higher than in the control group (Figures 5 B and 5C). The abundance of the genera Lysinibacillus, Bacteroides, and Acinetobacter in the healthy group is significantly higher than in the diseased group (Figure 5).

**Figure 5.**
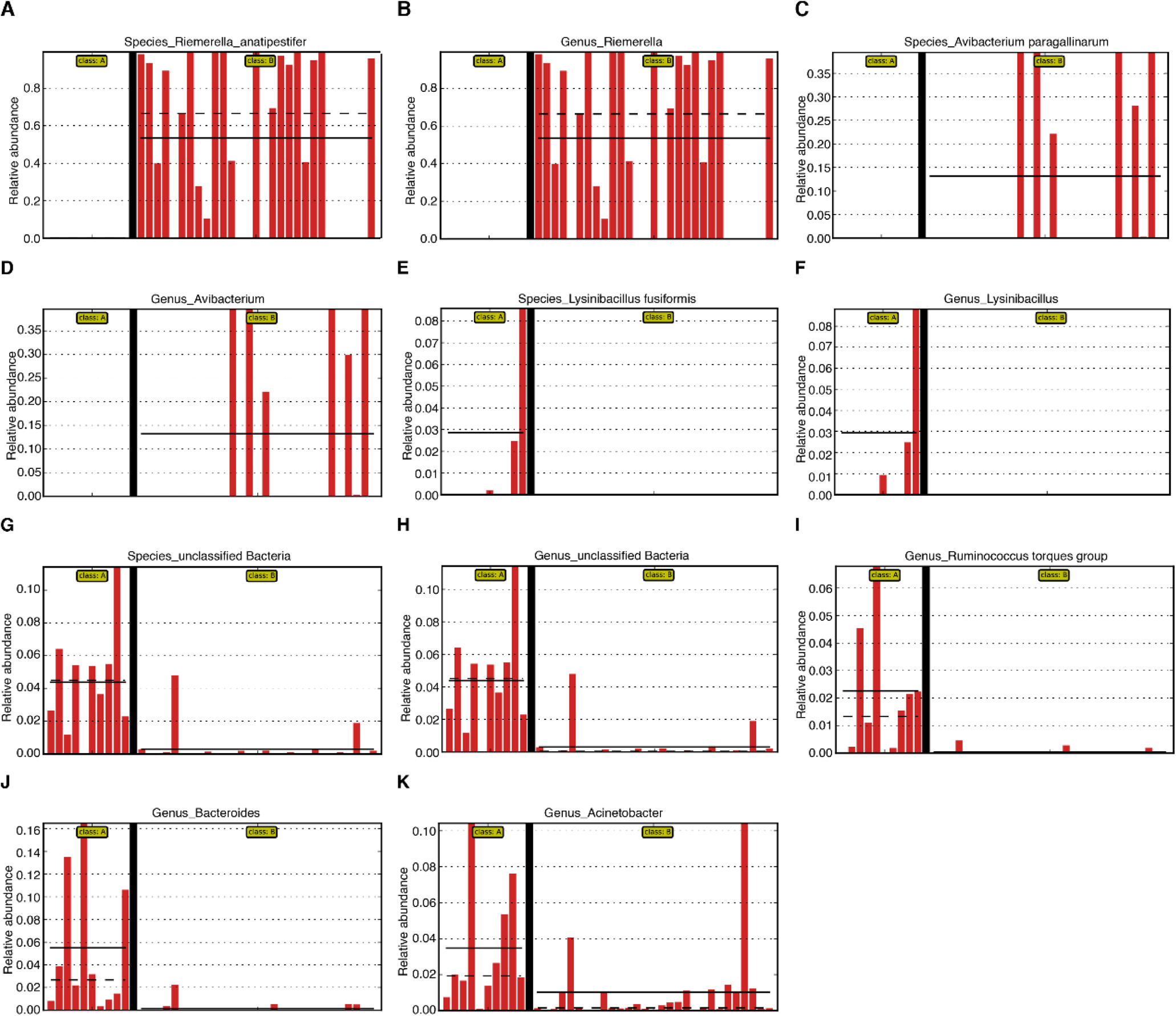
Intergroup abundance of biomarkers. For taxonomic units with significant differences between groups, LEfSe can also display their relative abundance distribution in different sample groups (identified by “class”), and use solid and dashed lines to indicate the mean and median relative abundance of the taxonomic unit in each group, respectively, thus visually demonstrating the differences between groups.

### 3.7 Functional gene prediction analysis

Functional gene prediction was performed using BugBase phenotype prediction (Figures 6 and 7). BugBase first normalizes OTUs by predicted 16S copy number, then predicts microbial phenotypes using the provided precalculated files. BugBase then selects the coverage threshold with the highest variance across all samples for each feature in the user’s data. The microbiota in the health and disease groups were compared using functional gene prediction. Overall, the relative abundance of aerobic, anaerobic, mobile-element-containing, facultatively anaerobic, biofilm-forming, gram-positive, potentially pathogenic, and stress-tolerant forms in the health group is higher than in the diseased group (Figures 6 and 7). The relative abundance of Gram-negative bacteria in the disease group is distinctly higher than that in the healthy group (Figure 6F). Aerobic and anaerobic bacteria are dominant in the healthy group (Figures 7A and 7B), whereas the disease group shows a higher relative abundance of facultatively anaerobic bacteria (Figure 7D).

**Figure 6.**
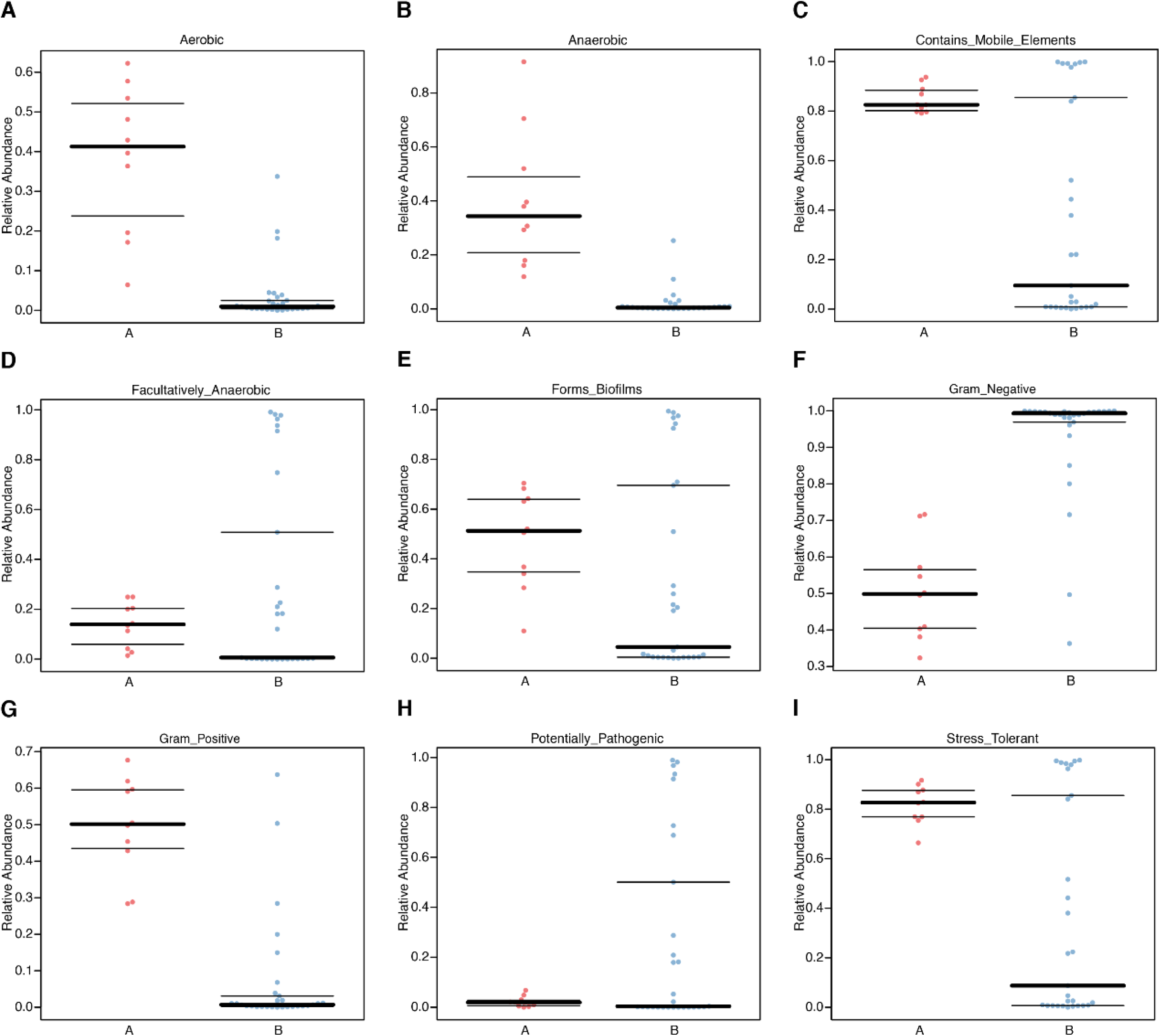
Functional gene prediction analysis. BugBase phenotypic prediction graph. The X axis represents the group name, and the Y axis represents the relative abundance percentage. The three lines, from bottom to top, represent the lower quartile, the mean, and the upper quartile, respectively.

**Figure 7.**
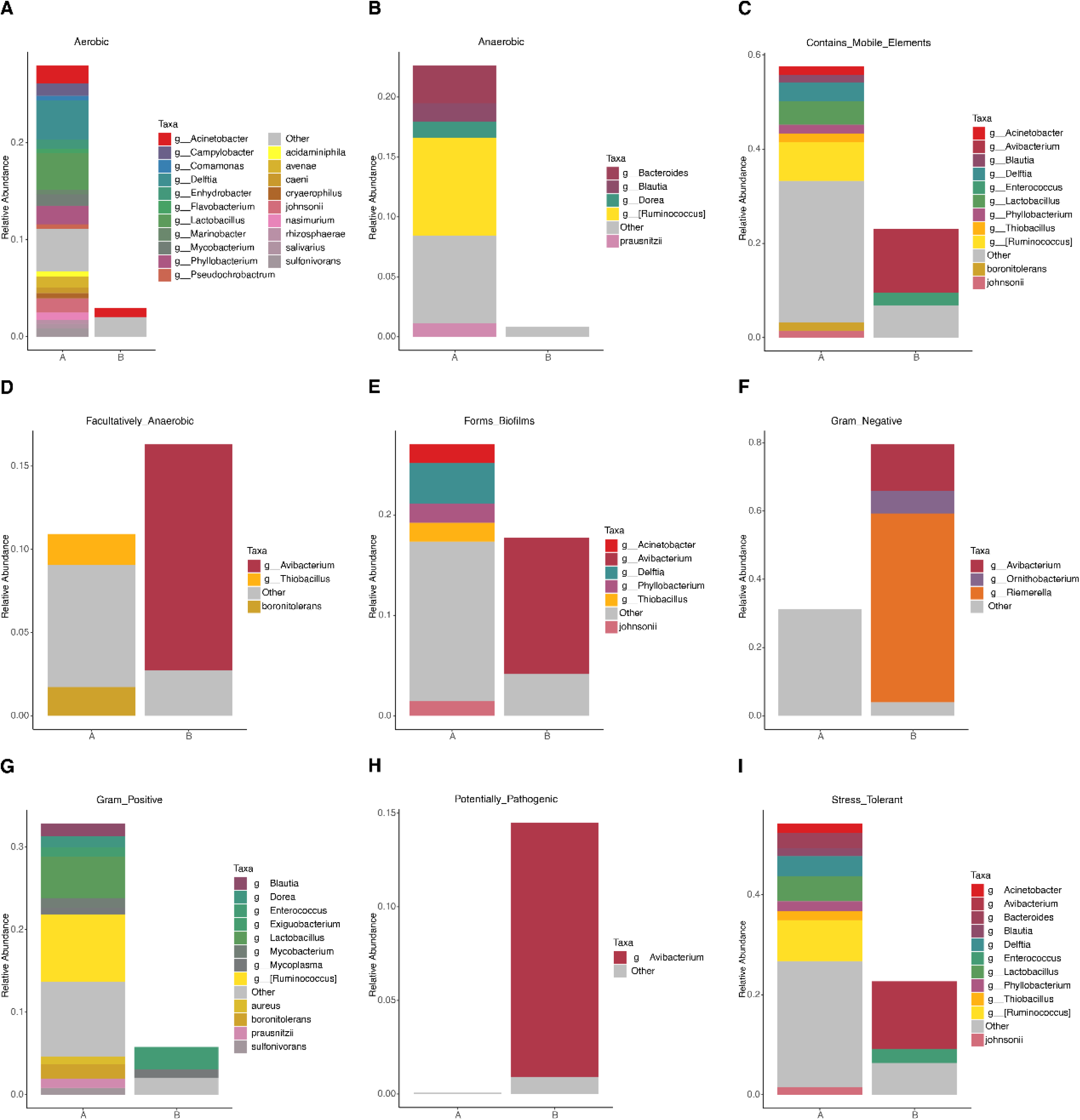
BugBase Species Bar Chart. The X axis represents the group name, and the Y axis represents the percentage of relative species abundance. The image shows nine phenotypes at the family level; additional taxonomic bar charts are available in the results file.

## 4 Discussion

Microbial communities are in symbiosis with the host, contributing to a balance between the host and the microbiota. The microbiota in humans and animals are crucial for the host and have been highlighted by numerous studies since their discovery ^20^. Genera *Escherichia*-*Shigella*, *Enterococcus*, *Staphylococcus,* and *Acinetobacter* were simultaneously present in six segments of the reproductive tract of laying hens, but with varied abundances. *Enterococcus*, *Gallicola*, *Corynebacterium*, and Escherichia-Shigella were the four most abundant genera in the cloaca. In contrast, Escherichia-Shigella, Enterococcus, Pseudomonas, Acinetobacter, and *Staphylococcus* represented the majority of the genera in the other five reproductive tract sections. Specifically, the potential pathogenic bacteria of Escherichia-Shigella exhibited the highest abundance in the segments from the Vagina to the infundibulum. A similar pattern was also observed in the genus *Pseudomonas* ^21^.

Environmental change and pathogen invasion may lead to dysbiosis of the microbiota. Microbiota dysbiosis is associated with infectious diseases, and microbiota surveys can be used to diagnose pathogenic bacteria ^22^. Pathogenic bacteria could be identified from the reproductive tract of laying hens, including *Escherichia-Shigella*, *Enterococcus*, *Staphylococcus*, and *Ornithobacterium* ^23^. In the present study, compared with the healthy control group, *Riemerella anatipestifer*, *Escherichia coli*, *Avibacterium paragallinarum*, *Ornithobacterium rhinotracheale*, and *Enterococcus faecalis* were identified at high relative abundance in the reproductive tract of the diseased group. *Arcobacter cryaerophilus* is a globally emerging foodborne and zoonotic pathogen ^17^. *E. faecalis* is significantly more associated with asymptomatic cases of primary endodontic infections than with symptomatic ones ^18^. Some bacteria, like *E. cecorum*, were initially known as symbiotic bacteria of chickens. Subsequent studies have shown that pathogenic strains of *E. cecorum* have become a significant cause of morbidity and mortality in broiler breeders, with repeated outbreaks reported ^19^. Synergetic infection with three isolates from diseased ducks was able to induce egg production, ruptured follicles, salpingitis, and saplings-peritonitis both in layers and breeder ducks ^24^. *Staphylococcus, Escherichia coli, Streptococcus*, and *Proteus* were also identified from the ovary of hens ^25^.

Pathogenic bacteria identified in the diseased animals are helpful for vaccine development and treatment ^26–28^. Vaccine candidates were identified and evaluated ^29^. The correlation of decreased *Salmonella* recovery with elevated lymphocyte and macrophage numbers strongly suggests that local cell-mediated immunity is involved in controlling SE injection in the ovaries and oviducts ^30^. The results indicate that PDAs could potentially be used to control SE colonization in the chicken reproductive tract; however, *in vivo* studies validating these results are warranted ^31^. Impacted oviduct constituted 0.87% of oviduct abnormality in commercial layer chicken with an overall mortality of 0.5%. The findings of this study showed that the oviduct impaction might be caused by E. coli in concurrence with *Proteous spp.*, *Klebsiella spp*., *Streptococcus spp*. and Mycoplasma ^32^.

Symbiotic bacteria have the potential to be used as dietary supplements for the prevention or treatment of disease. Dietary supplementation with probiotics can modulate the oviduct microbiota and improve the antioxidant status of laying hens, without causing tissue damage ^33^. *Lactobacillus crispatus* is effective against inflammation and has been studied for its role in treating or preventing oviductal inflammation. Supplementation with Lactobacillus crispatus supported ovarian health and follicle differentiation and preserved epithelial cell barrier function in the shell gland, reducing inflammatory damage in the genital tract. This dual efficacy underscores the probiotic’s role in lowering oviductal inflammation, regardless of the oviductal state ^34^. Transvaginal administration of *L. johnsonii* contributes to protection against infection in the oviduct by improving the microflora of the oviductal mucosa and strengthening the mechanical barrier function of the tight junctions ^35^. To prevent or treat the infectious disease, vaccine is the first choice to protect the host ^26^. *Bacteroides fragilis*, *Bacteroides salanitronis*, *Bacteroides barnesiae,* and *Clostridium leptum*, which were related to immune function and potentially contribute to enhanced egg production ^36^.

In conclusion, this study first analyzed the reproductive tract microbiota in health and disease by next-generation sequencing. *Riemerella anatipestifer*, *Escherichia coli*, *Avibacterium paragallinarum*, *Ornithobacterium rhinotracheale*, *Enterococcus faecalis*, and *Acinetobacter baumannii* were first identified as pathogenic bacteria at higher abundance in the disease group. Diseased group samples exhibited dysbacteriosis and a similar species-abundance cluster. Moreover, functional gene prediction analysis indicated that diseased groups showed a high level of gram-negative, facultative anaerobic, and potentially pathogenic bacteria. Next-generation sequencing is an effective method for analyzing bacterial communities to support health assessment and disease diagnosis.

## Acknowledgment

This study was funded by the Taishan Industrial Expert Programme (NO. tscx202306107). This study was supported by the Sichuan Natural Science Foundation (2025ZNSFSC1084).

## Author’s contribution

GF Zhang conceived and designed the study. Q Han, N Wang, T Feng, and YY Li executed the experiment. GF Zhang processed data analysis and writing. GF Zhang, KJ Quan, TL Xu, and Z Zhu edited the manuscript. All authors approved the final version of the manuscript.

## Data availability statement

The raw data supporting the conclusions of this article are available from the authors upon reasonable request.

